# Distinct patterns of selective sweep and polygenic adaptation

**DOI:** 10.1101/691840

**Authors:** Neda Barghi, Christian Schlötterer

## Abstract

The central paradigm of molecular population genetics is selective sweeps, where targets of selection have independent effects on the phenotype and quickly rise to fixation. In quantitative genetics, many loci contribute epistatically to adaptation and subtle frequency changes occur at many loci. Since both paradigms could result in a sweep-like genomic signature, additional criteria are needed to distinguish them. Using the framework of experimental evolution, we performed computer simulations to study the pattern of selected alleles under both paradigms. We identify several distinct patterns of selective sweeps and polygenic adaptation in populations of different sizes. These features could provide the foundation for development of quantitative approaches to differentiate the two paradigms.

**Author’s summary:** The selective sweep model assumes an independent frequency increase of favorable alleles and has been the basis of many tests for selection. While, polygenic adaptation is typically modelled by small frequency shifts in many loci. Recently, some theoretical and empirical work demonstrated that polygenic adaptation, similar to sweep, could also results in pronounced allele frequency changes. These results suggest that other distinct features need to be identified. Using computer simulations, we identified distinctive features for each paradigm that can be used to differentiate the sweep model from polygenic adaptation. Features such as allele frequency trajectories, time-series fitness, distribution of selected alleles on haplotypes, and parallelism among replicates can be used for development of suitable tests to distinguish between different adaptive architectures. These features provide the basis for theoretical modeling, design of selection experiments and data analysis.

## Introduction

Characterizing adaptive traits and, more recently, identification of their genetic basis has been one of the long-standing research fields in evolutionary biology. Molecular population genetic theory assumes that beneficial mutations are rare, but once occurred they rise in frequency until fixation (Smith and Haigh 1974), i.e. hard sweeps. More recently, the concept of classic hard sweeps have been extended - the beneficial allele either starts from standing genetic variation or multiple beneficial alleles are generated by mutation at the same gene (Hermisson & Pennings 2005). For decades the selective sweep paradigm has dominated molecular population genetics and the distortion of the allele frequency spectrum of sites flanking beneficial mutations has been exploited by a wealth of statistical tests to distinguish selection from neutrality (Messer and Petrov 2013; Pavlidis and Alachiotis 2017).

Quantitative genetics, on the other hand, traditionally has a strong focus on the evolution of phenotype, which is assumed to be determined by many contributing alleles, each with subtle effect, i.e. polygenic adaptation. Although adaptive traits are frequently assumed to be polygenic (Chevin and Hospital 2008; Pritchard and Di Rienzo 2010; Pritchard et al. 2010), the genomic signature of polygenic adaptation is being studied only recently. For polygenic adaptation small effect sizes are widely assumed to result in subtle allele frequency changes when a population is exposed to a new environment with a different trait optimum, trait optimum paradigm. Only recently, theoretical (Chevin and Hospital 2008; Höllinger et al. 2019; Jain and Stephan 2017) and empirical studies (Barghi et al. 2019) demonstrated that polygenic adaptation can also generate sweep-like selection signatures.

Sweep-like selection signatures arising from selective sweeps and polygenic adaptation suggests that the standard approach of studying genomic signatures in extant/evolved populations is not conclusive about the underlying paradigm. Because knowledge of the underlying paradigm of adaptation is crucial for the proper theoretical modelling and neutrality tests, alternative approaches are needed to distinguish between them and determine their importance for adaptation processes. Time series data provide information about the trajectories of beneficial alleles in evolving populations, which can be used to distinguish between the two paradigms. Time series data are, however, quite rare. In addition to fossil data, experimental evolution provides a powerful approach to study the adaptive architecture of traits (Kawecki et al. 2012; Schlotterer et al. 2015). The cost-effectiveness of sequencing pools of individuals (Schlöttereret al. 2014) provides the opportunity to generate time-series of genome-wide polymorphism data in multiple replicates.

Recently, the extent of genomic similarity among replicates was used as a summary statistic to determine the underlying evolutionary paradigm in 10 experimental replicates of *Drosophila simulans* (Barghi et al. 2019). With a single discriminating summary statistic not being powerful enough, in this study we aim to identify additional patterns in the evolving populations which are informative for recognizing the underlying evolutionary paradigm.

Reasoning that genetic drift provides a major perturbation of the directed forces of selection, we explored the potential of different experimental population sizes to distinguish between the paradigms. Using computer simulations, we identify several parameters such as allele frequency trajectories, time-series fitness, distribution of selected alleles on haplotypes, and parallelism among replicates, that distinguish sweep and trait optimum paradigms.

## Results and Discussion

With recent theoretical (Chevin and Hospital 2008; Höllinger et al. 2019; Jain and Stephan 2017) and empirical studies (Barghi et al. 2019) demonstrating that polygenic adaptation can also result in sweep-like selection signatures, is has become clear that the distinction of the underlying selection paradigm requires new approaches building on multiple diagnostic features. For example, we recently showed that evolutionary paradigms can be distinguished by the extent to which targets of selection are shared among replicates (Barghi et al. 2019). However, a reliable distinction between paradigms requires identification of additional features that distinguish both paradigms. We performed computer simulation under sweep and trait optimum paradigms with small and large populations sizes to identify distinct patterns for each paradigm. Our computer simulations are not designed to exhaustively cover all possible parameter combinations, but we rather identify distinct features of each paradigm.

### Distinct characteristics of sweep and trait optimum paradigms

We explored potential differences between selective sweep and trait optimum paradigms using a standard set of simulation parameters. In a population of 450 diploid individuals, 100 linked loci, matching typical E&R experiments in *Drosophila* (Barghi et al. 2019), with equal starting frequency of 0.05 and equal effects (selection coefficient of 0.08 for selective sweep and effect size of 0.04 for trait optimum paradigm) were simulated with the *D. simulans* recombination landscape in 500 iterations (scenario A in Table 1a, for sweep, and 1b, for trait optimum paradigm).

**Table 1.** Simulation parameters for sweep and trait optimum paradigms

| a. Sweep paradigm | A<br>default | B<br>different no. of loci | C<br>different s |
| --- | --- | --- | --- |
| Parameters |  |  |  |
| N | 450, 9000 | 450, 9000 | 450, 9000 |
| No. of loci | 100 | 10, 20, 50, 100 | 100 |
| Selection | 0.08 | 0.08 | 0.02, 0.05, 0.08, 0.1 |
| Starting frequency | 0.05 | 0.05 | 0.05 |
| Recombination map | D. simulans | D. simulans | D. simulans |
| b. Trait optimum paradigm | A<br>default | B<br>different no. of loci | C<br>different effect size |
| Parameters |  |  |  |
| N | 450, 9000 | 450, 9000 | 450, 9000 |
| No. of loci | 100 | 10, 20, 50, 100 | 100 |
| Effect size | 0.04 | 0.04 | 0.04, 0.08, 0.2, 0.4 |
| Fitness function | Guassian fitness function with standard deviation of 0.3, fitness ranges between 0.5 and 4.5. Optimum phenotype is -2.5 | Guassian fitness function with standard deviation of 0.3, fitness ranges between 0.5 and 4.5. Optimum phenotype varies depending on the no. of loci<sup>1</sup> | Guassian fitness function with standard deviation of 0.3, fitness ranges between 0.5 and 4.5. Optimum phenotype varies depending on the effect size of loci<sup>2</sup> |
| Starting frequency | 0.05 | 0.05 | 0.05 |
| Heritability | 0.5 | 0.5 | 0.5 |
| Recombination map | D. simulans | D. simulans | D. simulans |

<sup>1</sup> Fitness-phenotype functions are shown in Fig. S4a
<sup>2</sup> Fitness-phenotype functions are shown in Fig. S4b

Typical E&R studies have relatively small population sizes, which requires accounting for the expected allele frequency change (AFC) due to genetic drift to distinguish selection from neutrality. Regardless of the selection paradigm, genetic drift is quite strong in small populations (Fig. S1). We accounted for this by computing a frequency cut-off based on the 95% quantile of AFC under neutral simulations and only alleles with more extreme AFCs were considered to be selected (Fig. S1).

#### Allele frequency trajectorie

One important difference between the two paradigms is the pattern of allele frequency changes. Under the selective sweep paradigm, selected alleles continuously increase in frequency until they reach fixation (Fig. 1) whereas distinct phases of allele frequency changes were discerned for the trait optimum paradigm (Franssen et al. 2017). In the initial phase of adaptation, when the population is far from the trait optimum, most alleles increase in frequency (Fig. 1). After the phenotypic optimum is reached (generation 40, Fig. 2), the second phase starts where the allele frequencies plateau. However, drift affects this phase and in small populations this phase is either very short or not present at all. In small population, drift decreases the frequency of some alleles below the threshold for identification of selected alleles, and with loss of these alleles, the median frequency of the remaining alleles continues to rise (Fig. 1). The third phase of allele frequency changes includes fixation and loss of selected alleles. The first 2 phases are shown in Fig. 1; the third phase becomes noticeable after more generations e.g. 2500 (Fig. S2). We illustrate the first two phases of the trait optimum paradigm by showing the trajectories of alleles in a single replicate in Fig. S3.

**Figure 1.**
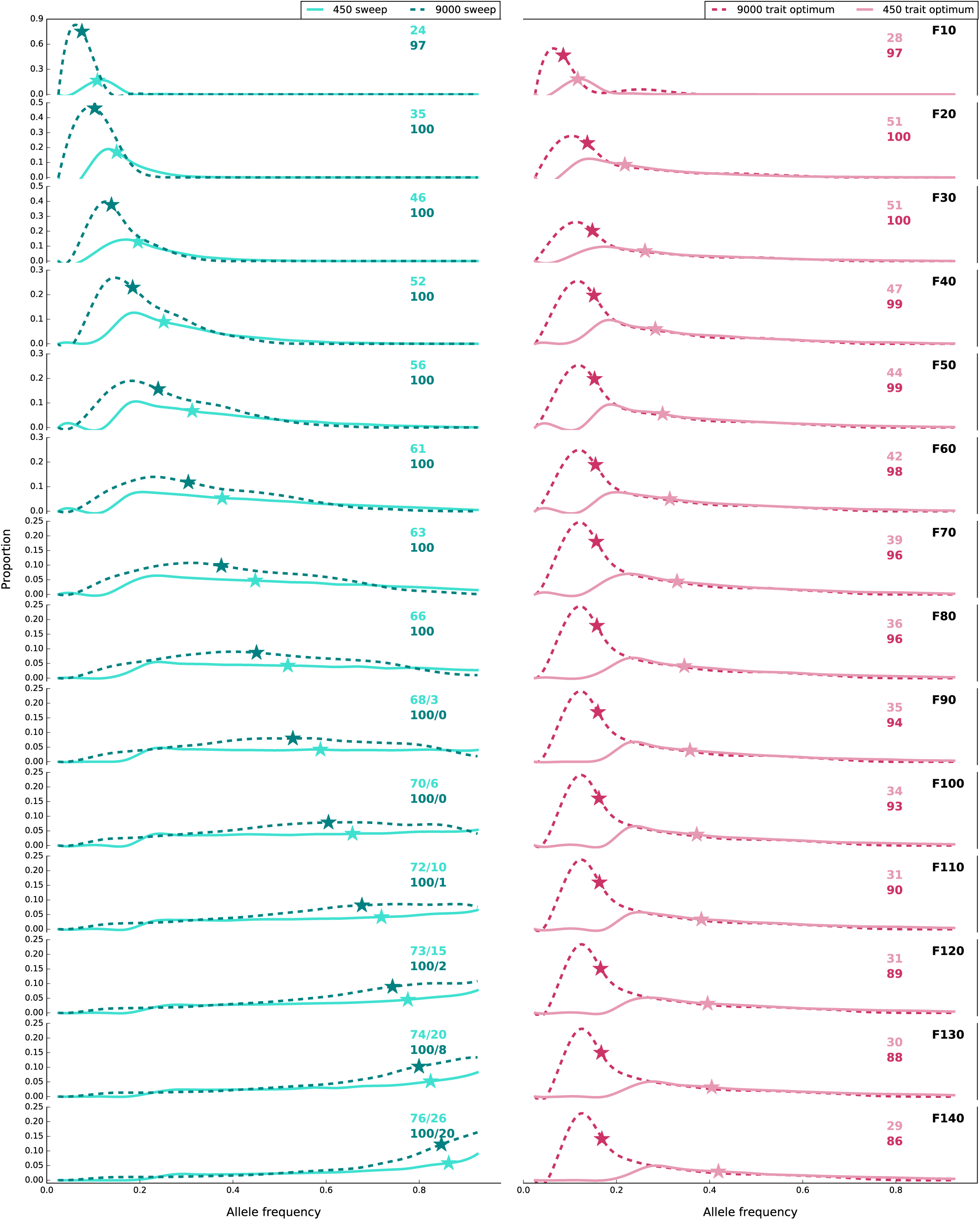
Frequency changes of alleles in populations of 450 and 9,000 individuals under sweep (left panel) and trait optimum (right panel) paradigms for scenario A (Table 1a and 1b). Asterisks depict the median frequency of alleles with frequency increase more than expected under drift. The number of alleles with frequency increase is shown with colors that correspond to the labels. The number of alleles with sweep-like signature (frequency ≥ 0.9), if present, is shown after ‘/’. The curves are fitted to histograms with bins of 0.05 and are normalized by bin count/total count. Total count is 50,000 (100 loci * 500 replicates).

**Figure 2.**
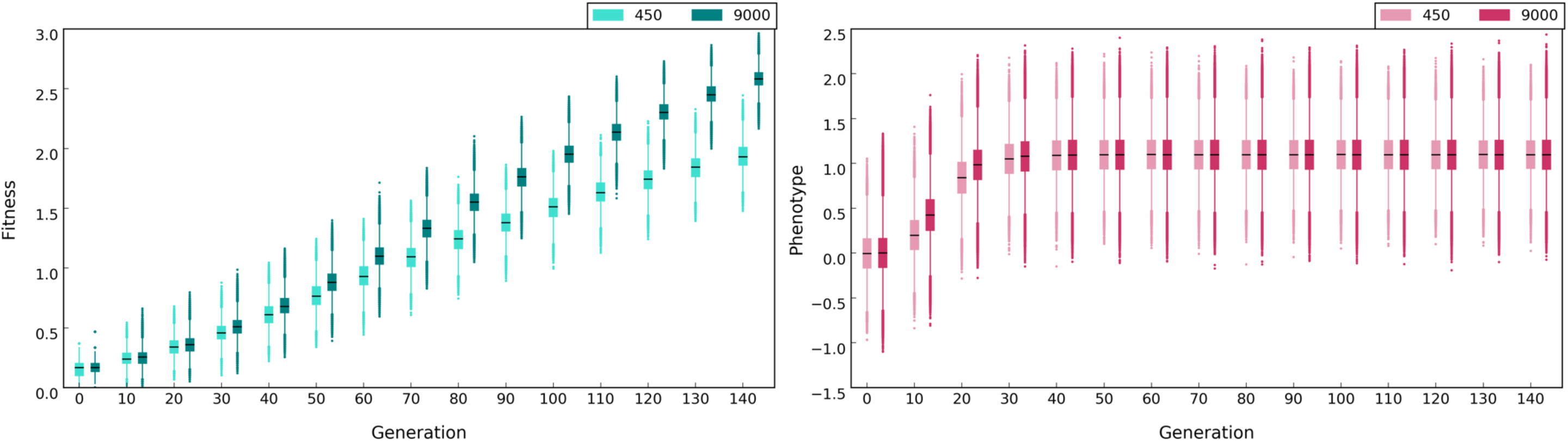
Population fitness and phenotype under sweep (left panel) and trait optimum paradigms (right panel) in populations of 450 and 9,000 individuals for scenario A (Table 1a and 1b). Black lines depict the median fitness or phenotype. Fitness is log_10_ transformed. The optimum phenotype in trait optimum paradigm is 1.1. The phenotype is normalized by subtracting the phenotype of each individual at F0 from the phenotype at each subsequent time point.

#### Sweep-like signatures

Many alleles reach frequency of ≥0.9, that is they exhibit sweep-like signatures, in the sweep paradigm (generation 90 onwards) while such signatures are not observed in the trait optimum paradigm (Fig. 1).

#### Fitness

As the frequency of selected alleles rises under the selective sweep paradigm (Fig. 1), population fitness also increases until all selected alleles are fixed (Fig. 2). Unlike the sweep paradigm, the population fitness under the trait optimum paradigm increases only until the phenotypic optimum is reached (Fig. 2). One distinct feature of the two paradigms is that for sweep paradigm the phenotypic value increases as long as the frequency of selected alleles do so. For the trait optimum paradigm, allele frequency changes are decoupled from the phenotype as soon as the trait optimum has been reached (Fig. 1 and 2).

#### Parallelism across replicates

Because the loss of alleles is more common in small populations due to drift, the different selected alleles may be detected among replicates resulting in lower parallelism among replicates (Fig. 3). This feature is shared between the two paradigms. For the sweep paradigm, parallelism continues to increases as more alleles reach frequencies above neutrality. In trait optimum paradigm, the contributing loci have epistasis for fitness and thus genetic redundancy is an intrinsic feature of the paradigm. Genetic redundancy describes the phenomenon that more alleles are segregating in a population than needed to reach the trait optimum (Barghi et al. 2019; Goldstein and Holsinger 1992; Nowak et al. 1997; Yeaman 2015). In this case, if some alleles contributing to the phenotype are lost, the trait optimum can still be reached by frequency increase of the remaining alleles. In the trait optimum paradigm, parallelism increases until populations reach the phenotypic optimum but it decreases afterwards (Fig. 3). This pattern can be explained by some alleles decreasing their frequency below the detection cutoff (Fig. 1). Since the stochasticity of the small populations in the first phase results in different loci contributing to the reach of trait optimum, the loss of alleles due to stochasticity in the second phase reduces the parallelism even more (Fig. 3).

**Figure 3.**
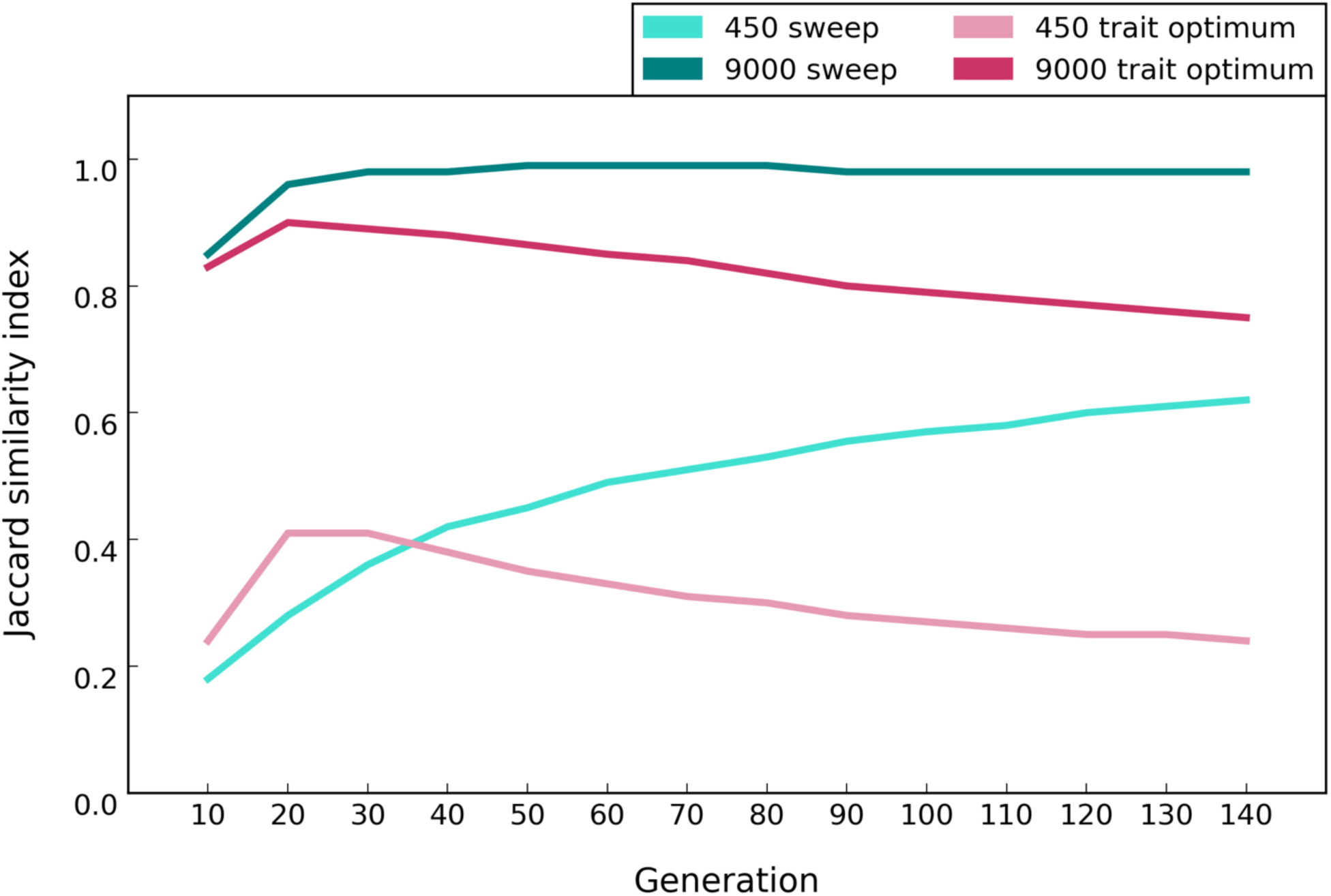
Median Jaccard similarity index in populations of 450 and 9,000 individuals under sweep and trait optimum paradigms for scenario A (Table 1a and 1b). Jaccard index for 10 replicate populations quantifies the extent to which alleles are shared among replicates (0 = no overlap, 1 = complete sharing). The average Jaccard index among replicates for 50 sets of 10-replicate evolution experiments were computed. For the trait optimum paradigm, the optimum phenotype is reached at generation 40 and 30 in small and large populations, respectively.

#### Distribution of selected alleles on haplotypes

In our simulations, the beneficial alleles were randomly distributed across the chromosomes in the founder populations so that each haplotype carries on average 5-6 beneficial alleles (Fig. 4). Due to recombination, the number of beneficial alleles per haplotype increases in both paradigms. While under the sweep paradigm, the number of beneficial loci per haplotype continues to increase (Fig. 4), for the trait optimum paradigm, this number increases only until the fitness optimum is reached (at F40) but does not change afterwards. Thus, another distinctive pattern between the two paradigms is the plateau in the number of beneficial alleles per haplotype in the trait optimum paradigm while this number continuously increases under the sweep paradigm until all alleles are fixed.

**Figure 4.**
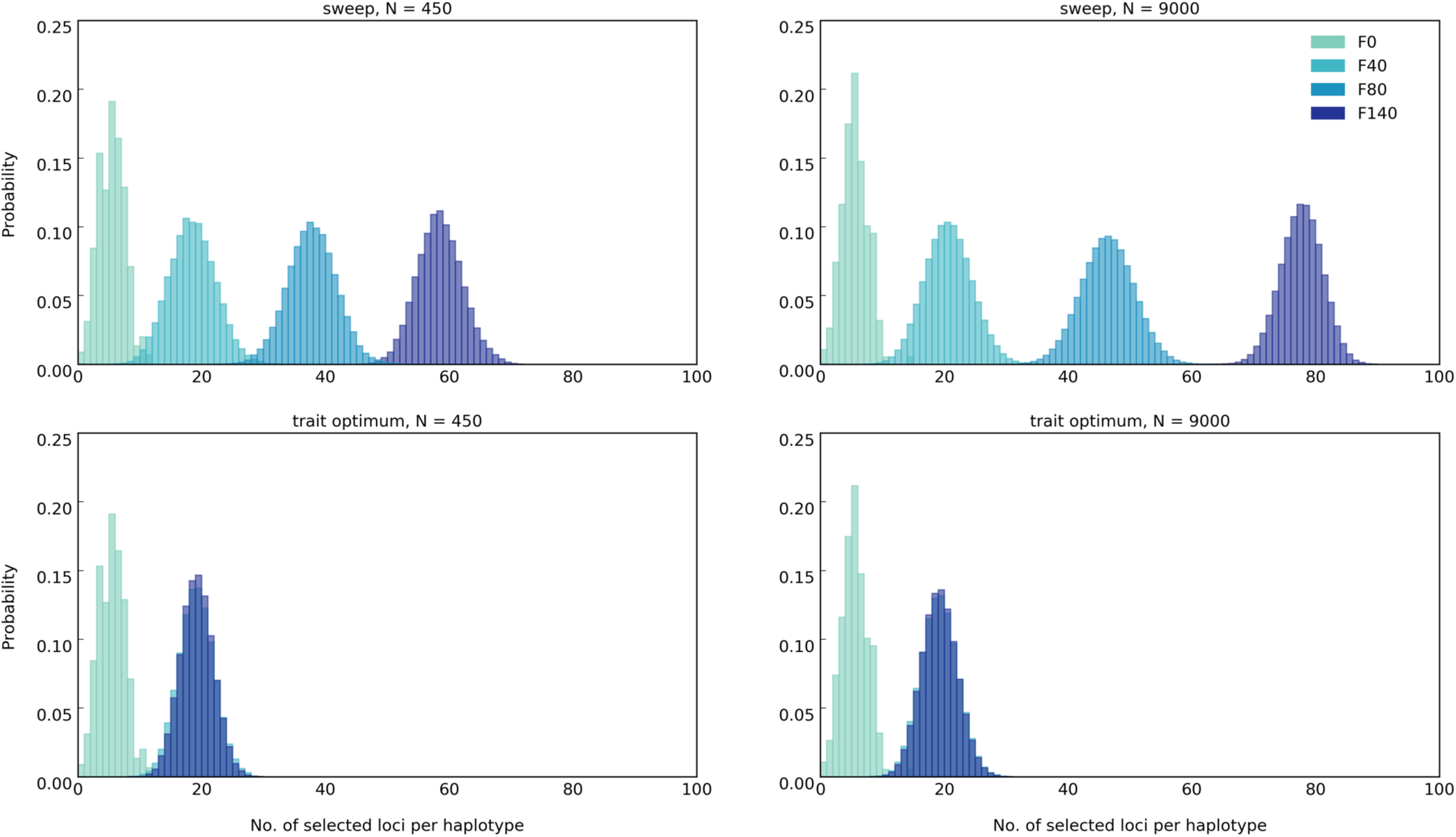
Number of beneficial loci in each haplotype under sweep (top panels) and trait optimum (bottom panels) paradigms for scenario A (Table 1a and 1b). The area under normalized histogram equals 1. Total number of replicates in simulations are 50.

### Effect of population size

Large populations experience less genetic drift than small ones, which increases the efficacy of selection and the power to detect selected alleles. To assess the impact of population size on the ability to discriminate between sweep and trait optimum paradigms, we also performed simulations with a larger population size, i.e. 9000 diploid individuals (scenario A in Table 1a, for sweep, and 1b, for trait optimum paradigm). Comparison of the sweep and trait optimum paradigms in small and large populations revealed additional distinctive features to differ between the paradigms possible only by the combined analysis of different population sizes.

#### Allele frequency changes

The neutral AFC in the large population is only 0.034 until generation 140, much less than in small populations (0.18, Fig. S1). Therefore, a plateau of the median allele frequencies after reaching the optimal trait under the trait optimum paradigm is observed in large populations (Fig. 1) which provides an unambiguous signature differentiating the two paradigms.

The difference in median allele frequencies between small and large populations increases with time for the trait optimum paradigm (Fig. 1). This pattern is the consequence of more loci decreasing below the detection limit in the small populations than for large ones after the trait optimum has been reached. For the sweep paradigm, the allele frequencies continuously increase with time so the difference in the median allele frequencies between small and large populations decreases continuously. Hence, large and small populations have characteristic signatures that distinguish trait optimum paradigm from sweep paradigm. Combining the information from large and small populations provides an even stronger distinction between the two paradigms.

#### Fitness

The evolution of fitness has the same trend in small and large populations regardless of the evolutionary paradigm. The increase in fitness is higher in large populations than in the small ones in the sweep paradigm (Fig. 2) because fewer alleles are lost by drift (Fig. 1). Furthermore, the population fitness increases faster in the large population under the trait optimum paradigm but only until the phenotypic optimum is reached (Fig. 2). Despite faster increase of fitness in large populations under trait optimum paradigm, small and large populations reach the fitness optimum almost at the same time (Fig. 2) and the differences in fitness between small and large populations before reaching the fitness optimum are very subtle. However, in the sweep paradigm, the difference in fitness gain between small and large populations increases with time. Thus, the differential fitness in populations of different sizes can serve as discriminator between the two paradigms.

#### Parallelism across replicates

Regardless of the evolutionary paradigm, the signature of selection is more repeatable in large populations than in small ones because fewer alleles are lost due to drift (Fig. 3).

#### Distribution of selected alleles on haplotypes

Under the sweep paradigm, more selected alleles are recombined onto the same haplotype in large populations than in small ones (Fig. 4) and the number of selected alleles increases with time for both population sizes. For the trait optimum paradigm, population size has no influence on the number of selected alleles on haplotypes after trait optimum is reached.

While for some discriminatory features, such as distribution of selected alleles on haplotypes, no major difference can be noted between large and small populations, contrasting the patterns of fitness evolution, allele frequency changes and parallelism among replicates in small and large populations clearly provides some additional information not available from analysis of a single population size alone.

### Effect of the number of selection targets

We determined the influence of the number of selected alleles by simulating 10, 20, 50 and 100 linked loci (scenario B in Table 1a, for sweep, and 1b, for trait optimum paradigm) with starting frequency of 0.05 and equal effects (0.08 for selective sweep and 0.04 for trait optimum paradigm) in small (450) and large (9000) populations in 500 iterations.

In the sweep paradigm, fitness of populations with more selected alleles is greater than that of populations with fewer alleles (Fig. 5) due to the frequency increase of more selected alleles throughout the time (Fig. 6). For the trait optimum we noticed a marked difference for founder populations with few alleles (e.g. 10 and 20), as in these simulations the trait optimum could not be reached (Fig. 7), hence no genetic redundancy was observed. In populations without redundancy (e.g. with 10 and 20 loci) the trajectories of allele frequencies (Fig. 8) resemble the sweep paradigm in that the median allele frequency continues to increase and the two paradigms cannot be distinguished.

**Figure 5.**
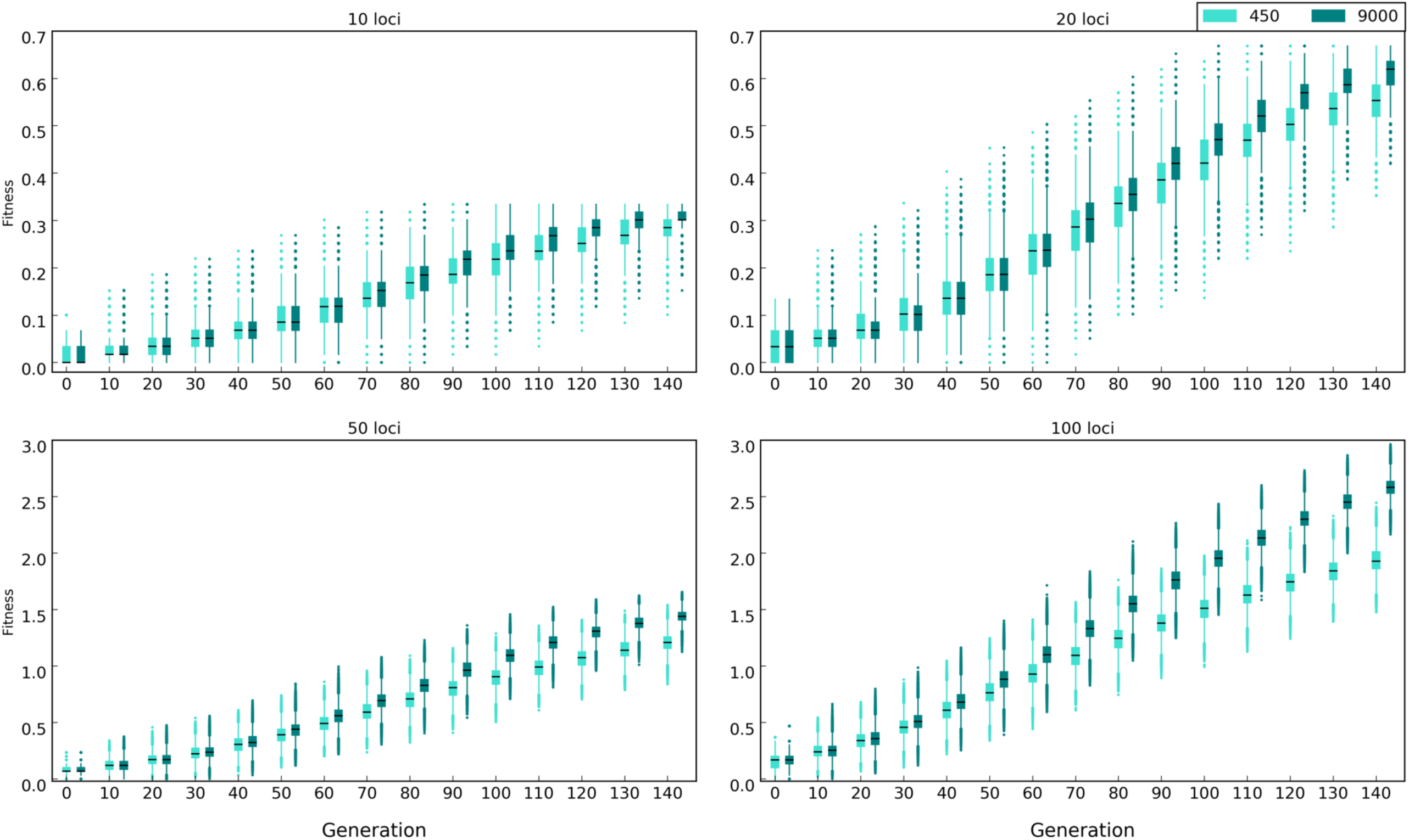
Population fitness with different number of beneficial loci, e.g. 10, 20, 50, and 100, under sweep paradigm (Table 1a, scenario B). Black lines depict the median fitness. Fitness is log_10_ transformed.

**Figure 6.**
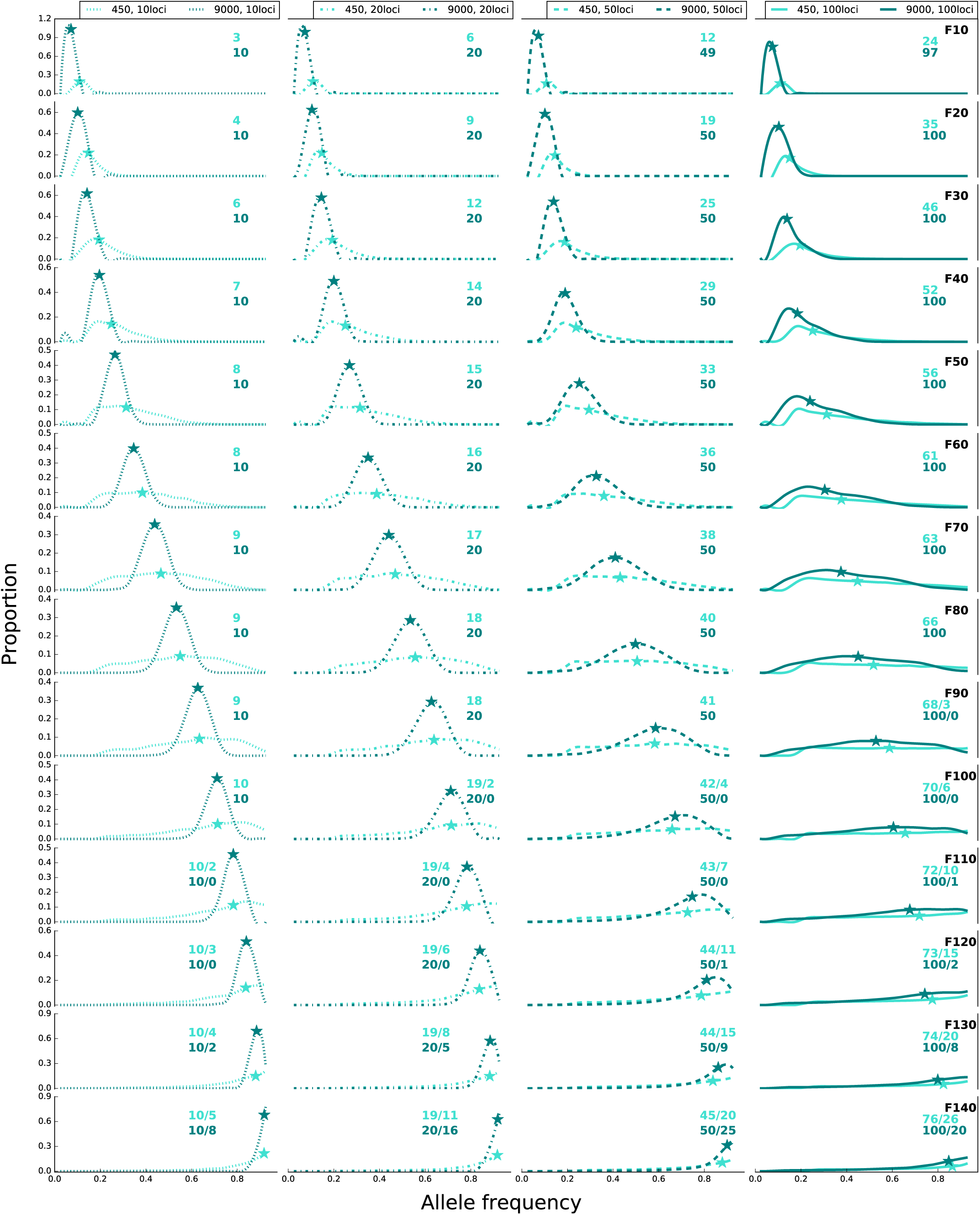
Frequency of alleles in populations of 450 and 9,000 individuals under sweep paradigm (Table 1a, scenario B) with different number of beneficial loci: 10 (dotted line), 20 (dash dotted line), 50 (dashed line), and 100 (solid line). Asterisks depict the median frequency of alleles with frequency increase more than expected under drift. The number of alleles with frequency increase is shown with colors that correspond to the labels. The number of alleles with sweep-like signature (frequency ≥ 0.9), if present, is shown after ‘/’. The curves are fitted to histograms with bins of 0.05 and are normalized by bin count/total count. Total count is number of loci * 500 replicates.

**Figure 7.**
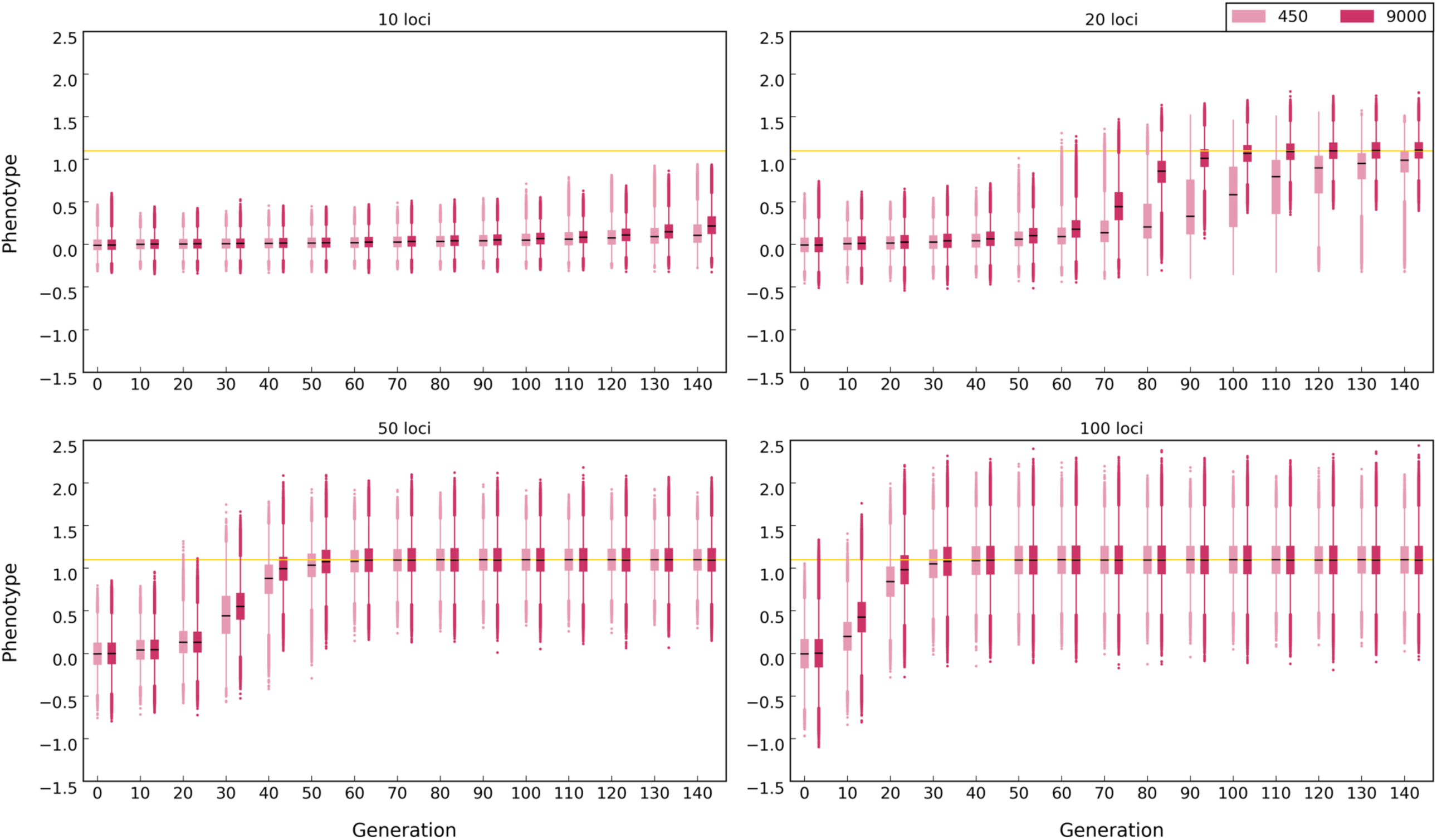
Phenotype in populations with 450 and 9,000 individuals with different number of beneficial loci under trait optimum paradigm (Table 1b, scenario B). Black lines depict the median phenotype. Yellow line shows the optimum phenotype. Note that the distance between the population phenotype at generation 0 and the optimum phenotype is equal across all simulations. The optimum phenotype is 1.1 (shown by yellow line). To normalize the phenotypes among different simulations, the phenotype of each individual at F0 is subtracted from the phenotype at each subsequent time point.

**Figure 8.**
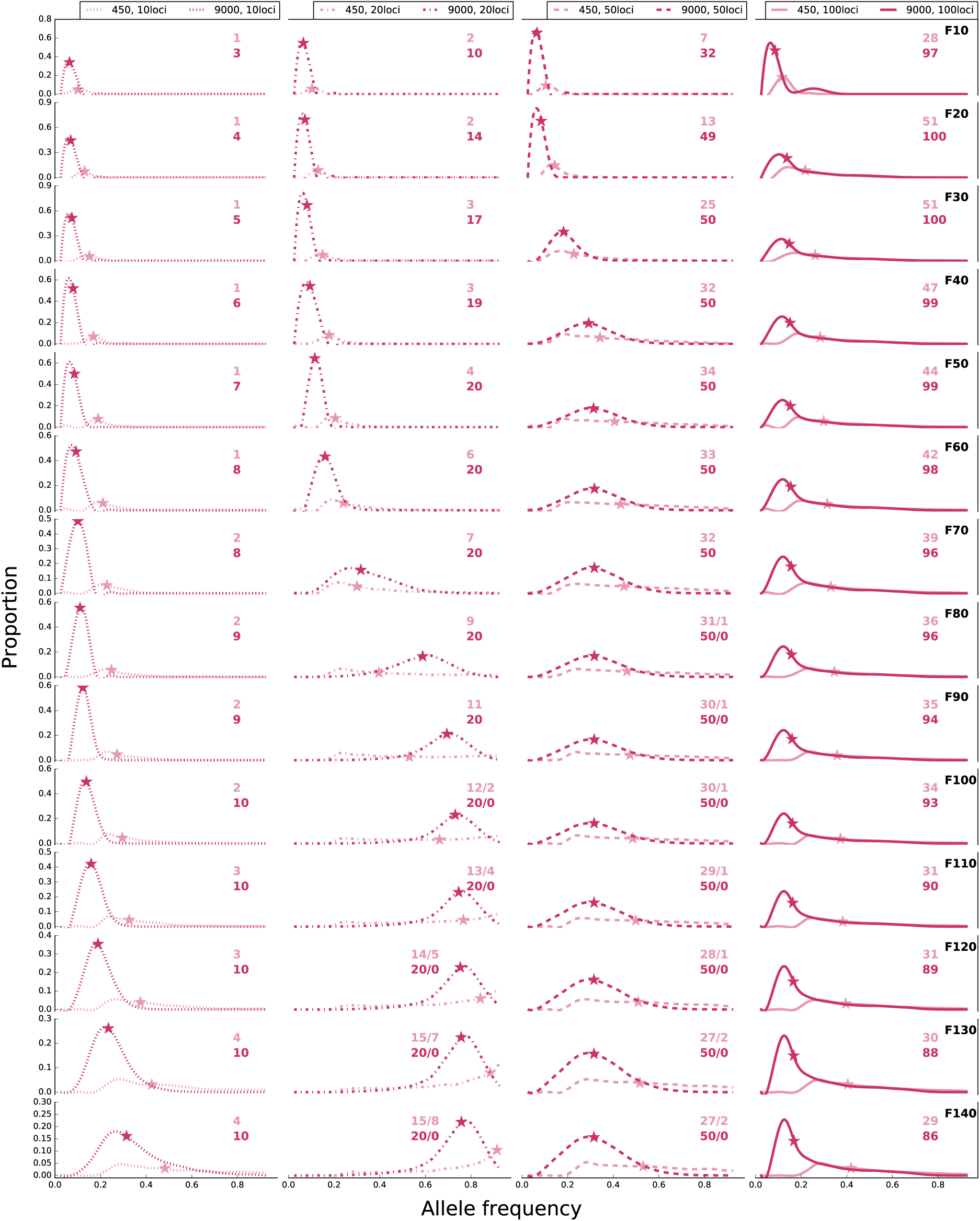
Frequency of alleles in populations of 450 and 9,000 individuals under trait optimum paradigm with different number of beneficial loci (scenario B in Table 1b): 10 (dotted line), 20 (dash dotted line), 50 (dashed line), and 100 (solid line). Asterisks depict the median frequency of alleles with frequency increase more than expected under drift. The number of alleles with frequency increase is shown with colors that correspond to the labels. The number of alleles with sweep-like signature (frequency ≥ 0.9), if present, is shown after ‘/’. The curves are fitted to histograms with bins of 0.05 and are normalized by bin count/total count. Total count is number of loci * 500 replicates.

Nevertheless, we noticed an interesting pattern: under the sweep paradigm in small populations the fraction of alleles with frequency change more than expected under neutrality decreases with the increase in the number of selection targets (Fig. 6). As shown previously (Barton 1995), this pattern is the outcome of selection at other loci causing variation in fitness. Recombination generates haplotypes with a larger variance in the number of selected alleles on a single haplotype. This in turn increases the variance in fitness for those populations, ultimately leading to the loss of some selected alleles by genetic drift. As a consequence, the similarity among replicates in populations with fewer selected alleles is greater than those with more alleles (Fig. 9).

**Figure 9.**
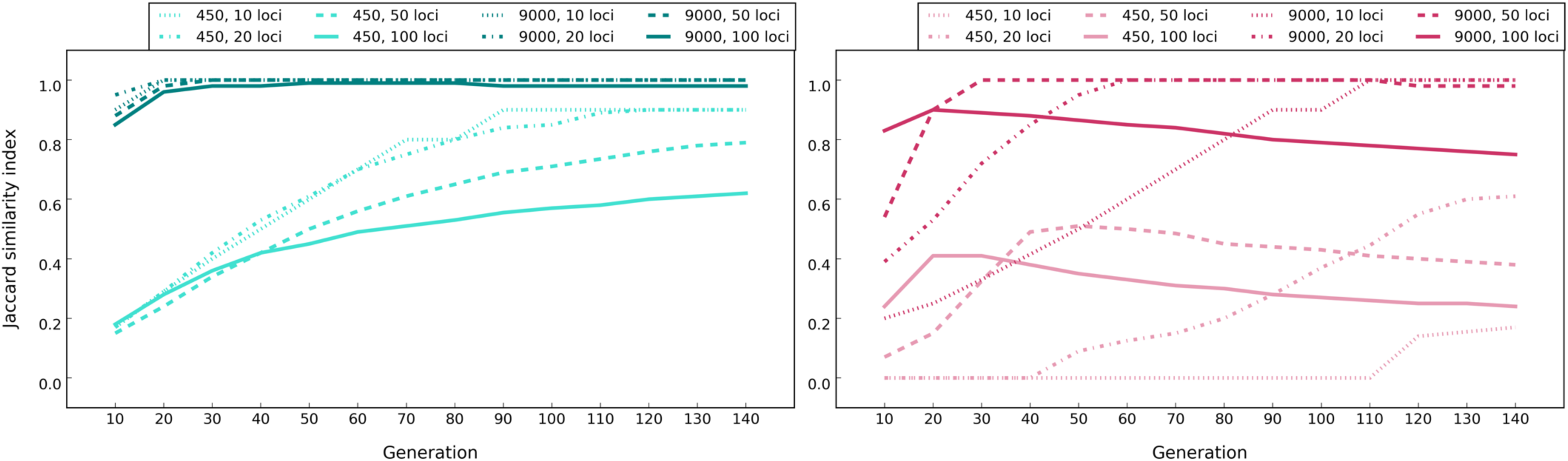
Median Jaccard similarity index in populations of 450 and 9,000 individuals with different number of beneficial loci under sweep and trait optimum paradigms simulations (scenario B in Table 1a and 1b). The average Jaccard index among replicates for 50 sets of 10-replicate evolution experiments were computed. For the trait optimum paradigm, the generation the optimum phenotype is reached is shown in Figure 7.

### Importance of allelic effect size

We evaluated the influence of allelic effect size/selection coefficient by simulating 100 linked loci with starting frequency of 0.05 and varying effect sizes (scenario C in Table 1a, for sweep, and 1b, for trait optimum paradigm) in small and large populations. Effect size/selection coefficient have pronounced influence on the evolutionary trajectories. Fitness increases faster with higher selection coefficients (Fig. 10) and sweep signatures (Fig. 11) become more frequent. The trait optimum is also reached faster with alleles of larger effect sizes (Fig. 12) because subtle shifts in frequency cause the required phenotypic shift (Fig. 13). Thus, the signatures of reaching trait optimum - drift reduces parallelism among replicates (Fig. 14) and the number beneficial alleles per haplotypes remains stable – are seen earlier (Fig. 15). Except for minor differences, the main distinctions between sweep and trait optimum paradigms do not change with different selection coefficients/effect sizes. For example, with small *s* (0.02) the median frequency of the selected alleles continues to increase faster in small populations than the subtle changes in the large ones (Fig. 11), similar to the trait optimum paradigm (Fig. 13). Nevertheless, for the sweep paradigm, the number of identified selection targets, even for small *s*, increases as populations evolve (Fig. 11), while it decreases for the trait optimum paradigm (Fig. 13). These distinctive patterns can be used for differentiating the paradigms.

**Figure 10.**
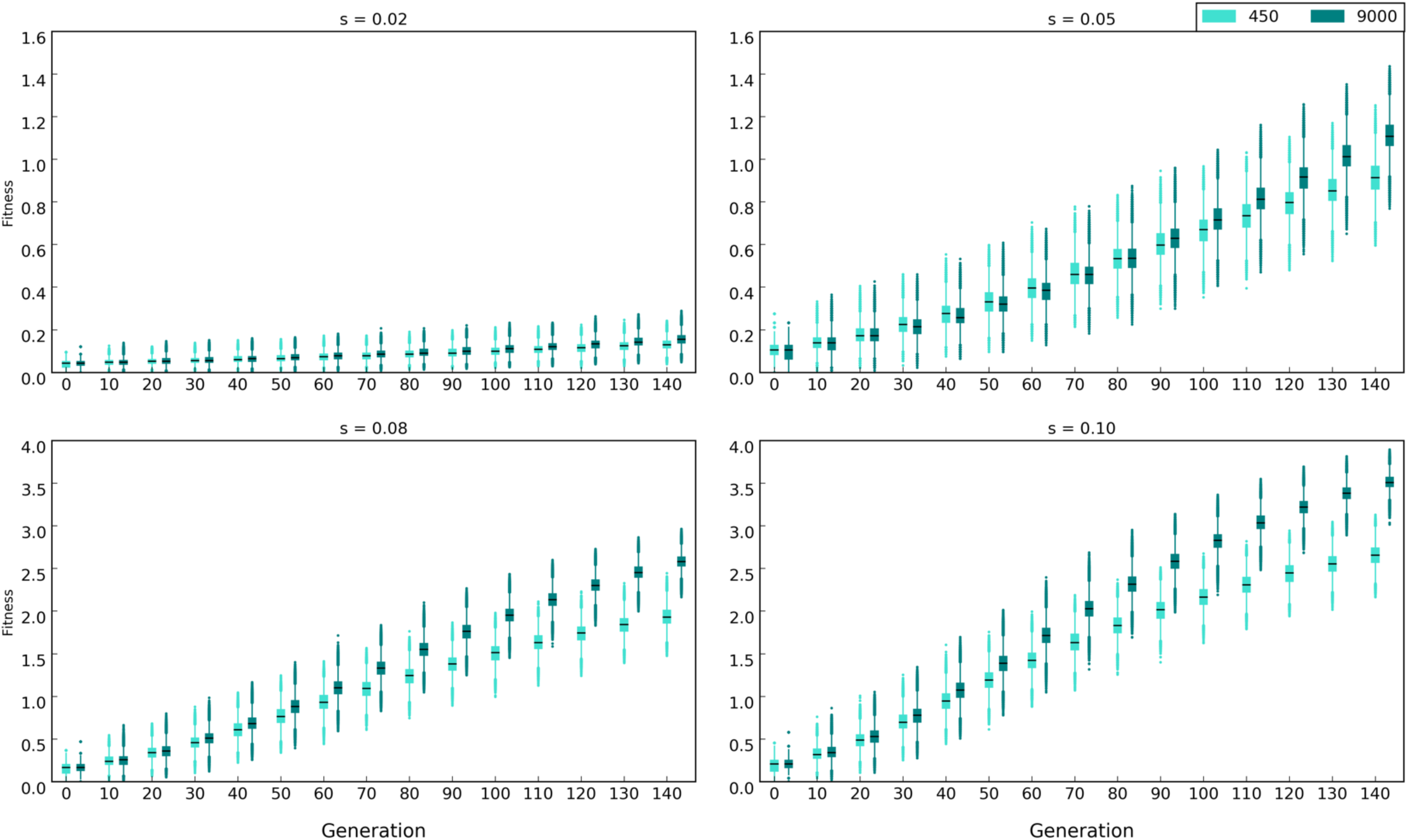
Fitness in populations of 450 and 9,000 individuals with 100 beneficial alleles and different selection coefficients (0.02, 0.05, 0.08 and 0.1) under sweep paradigm (Scenario C in Table 1a). Black lines depict median fitness. Fitness is log10 transformed

**Figure 11.**
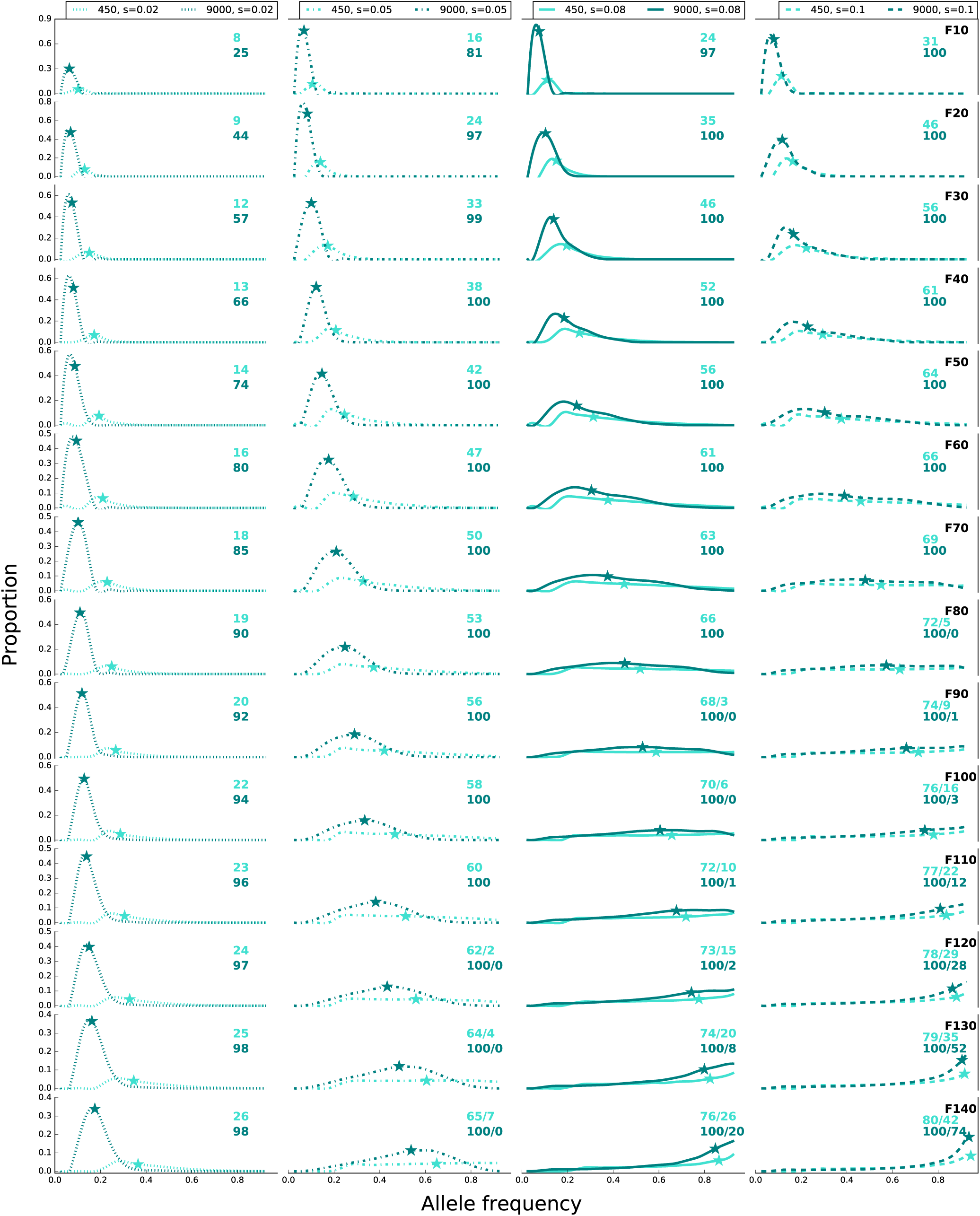
Frequency of alleles in populations of 450 and 9,000 individuals under sweep paradigm with 100 beneficial loci and different selection coefficients (scenario C in Table 1a): 0.02 (dotted line), 0.05 (dash dotted line), 0.08 (solid line), and 0.1 (dashed line). Asterisks depict the median frequency of alleles with frequency increase more than expected under drift. The number of alleles with frequency increase is shown with colors that correspond to the labels. The number of alleles with sweep-like signature (frequency ≥ 0.9), if present, is shown after ‘/’. The curves are fitted to histograms with bins of 0.05 and are normalized by bin count/total count. Total count is 100 loci * 500 replicates.

**Figure 12.**
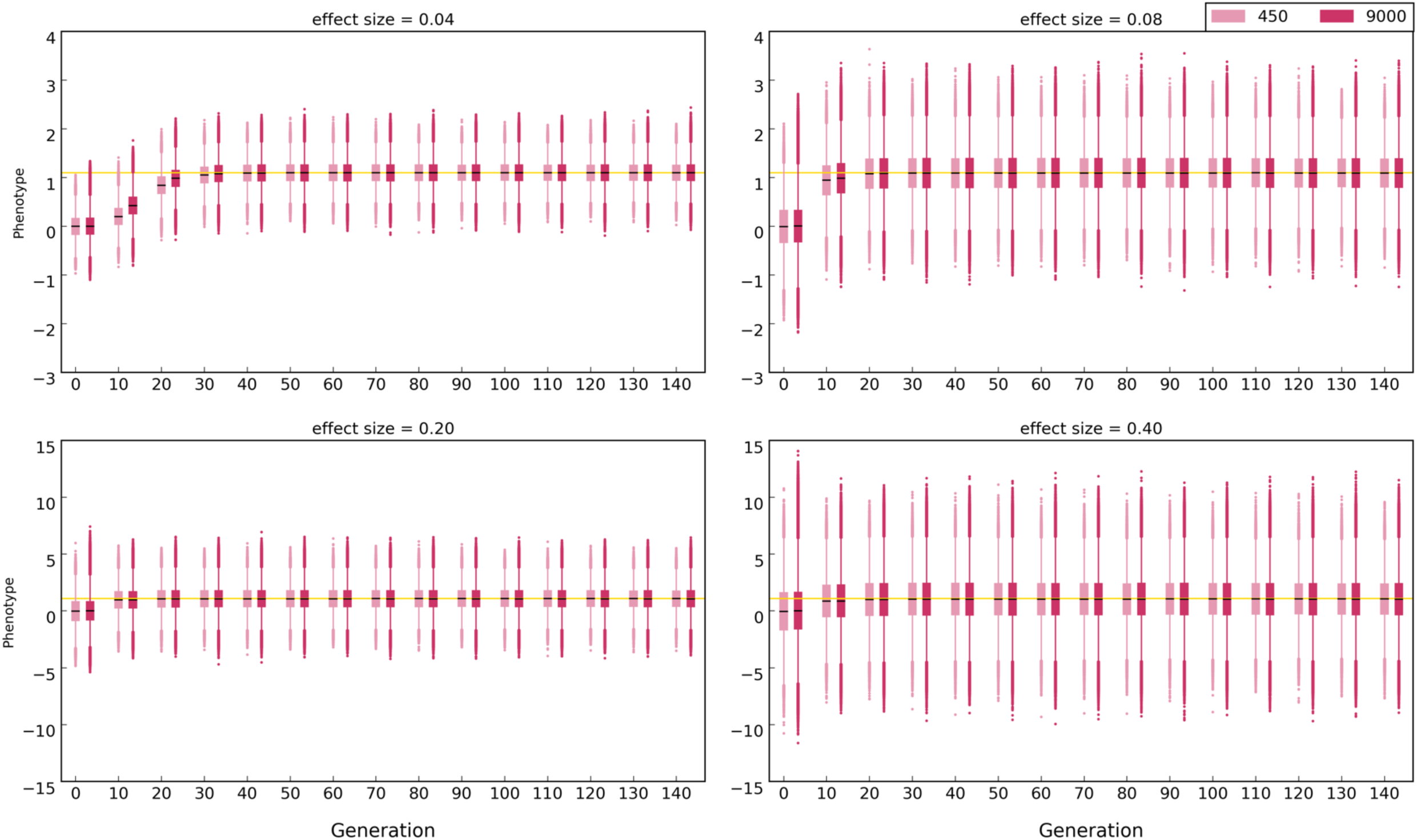
Phenotype in populations of 450 and 9,000 individuals with 100 beneficial loci and different effect sizes (0.04, 0.08, 0.2, 0.4) under trait optimum paradigm (scenario C in Table 1b). Black lines depict the median phenotype. Yellow line shows the optimum phenotype (1.1). Note that the distance between the population phenotype at generation 0 and the optimum phenotype is equal across all simulations. To normalize the phenotypes among different simulations, the phenotype of each individual at F0 is subtracted from the phenotype of at each subsequent time point.

**Figure 13.**
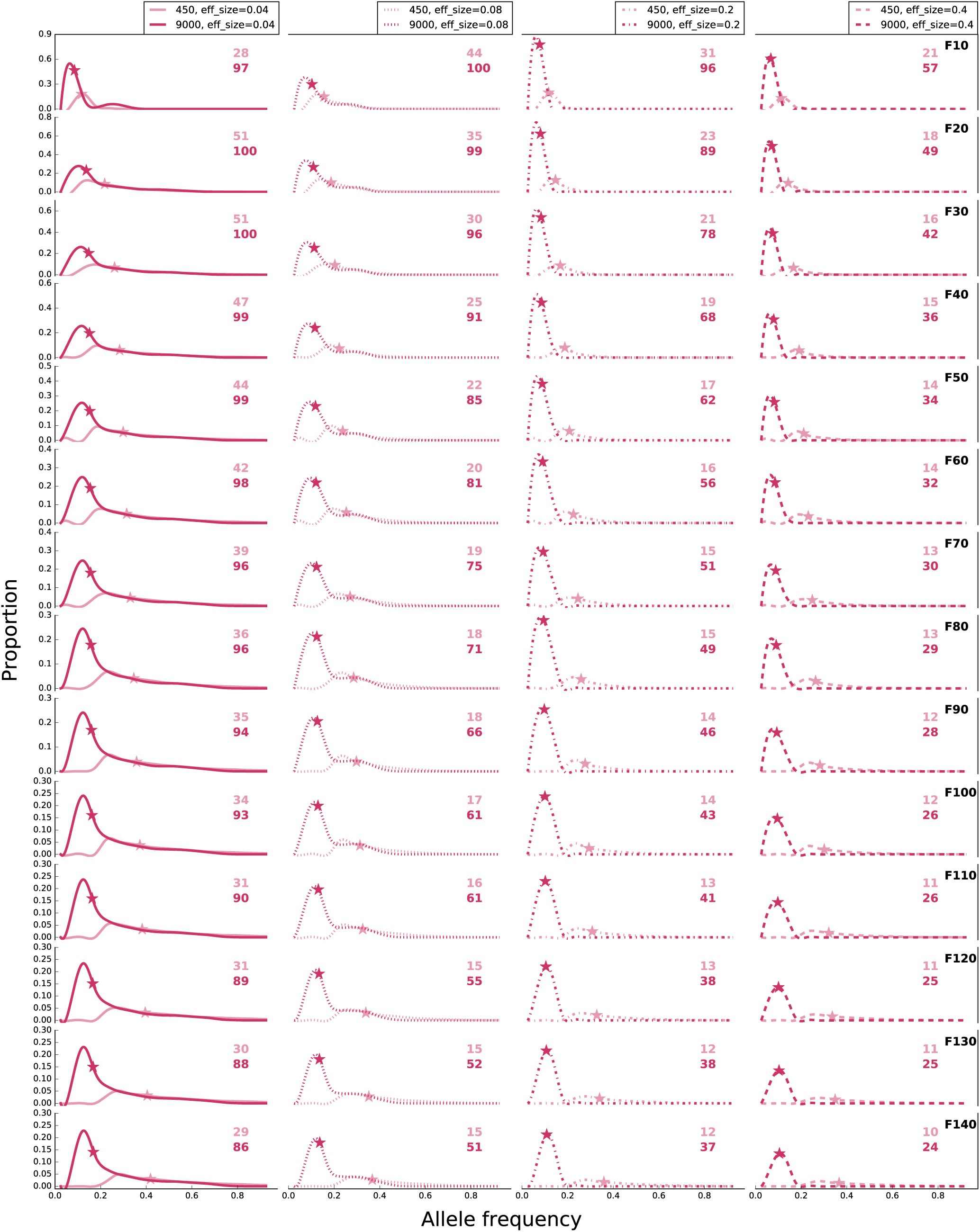
Frequency of alleles in populations of 450 and 9,000 individuals under trait optimum paradigm with 100 beneficial loci and different effect sizes (scenario C in Table 1b): 0.04 (solid line), 0.08 (dotted line), 0.2 (dashed dotted line), and 0.4 (dashed line). Asterisks depict the median frequency of alleles with frequency increase more than expected under drift. The number of alleles with frequency increase is shown with colors that correspond to the labels. The number of alleles with sweep-like signature (frequency ≥ 0.9), if present, is shown after ‘/’. The curves are fitted to histograms with bins of 0.05 and are normalized by bin count/total count. Total count is 100 loci * 500 replicates.

**Figure 14.**
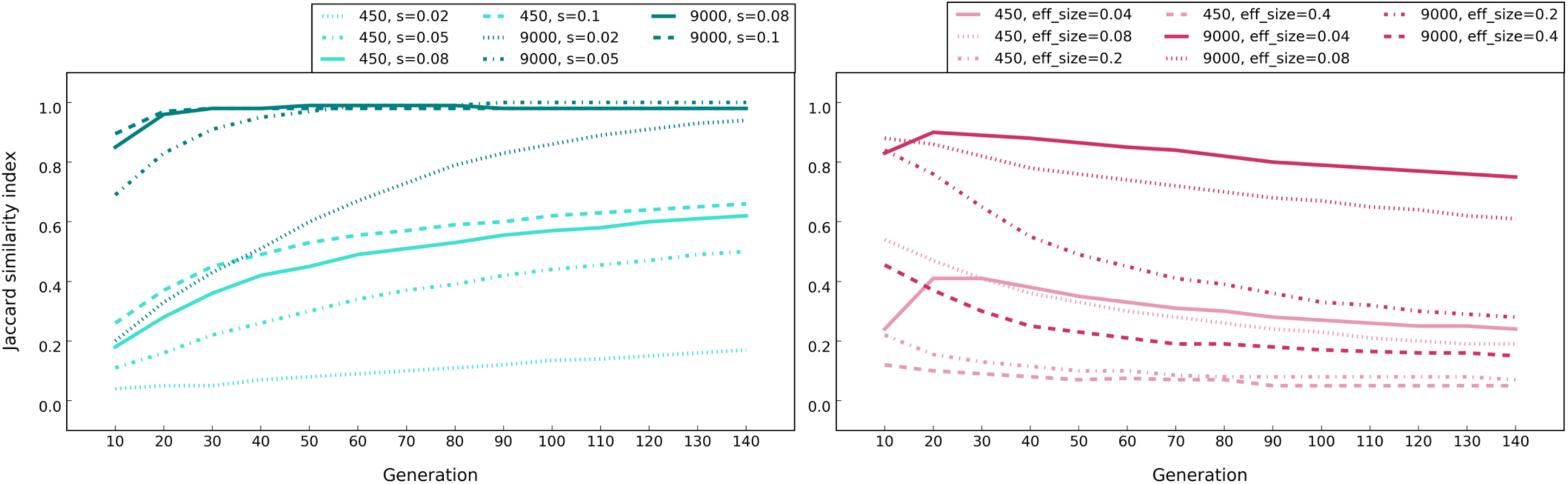
Median Jaccard similarity index in populations of 450 and 9,000 individuals with 100 loci and different effect sizes under sweep and trait optimum paradigms (scenario C in Table 1a and 1b). The average Jaccard index among replicates for 50 sets of 10-replicate evolution experiments were computed. For the trait optimum paradigm, the generation the optimum phenotype is reached is shown in Figure 12.

**Figure 15.**
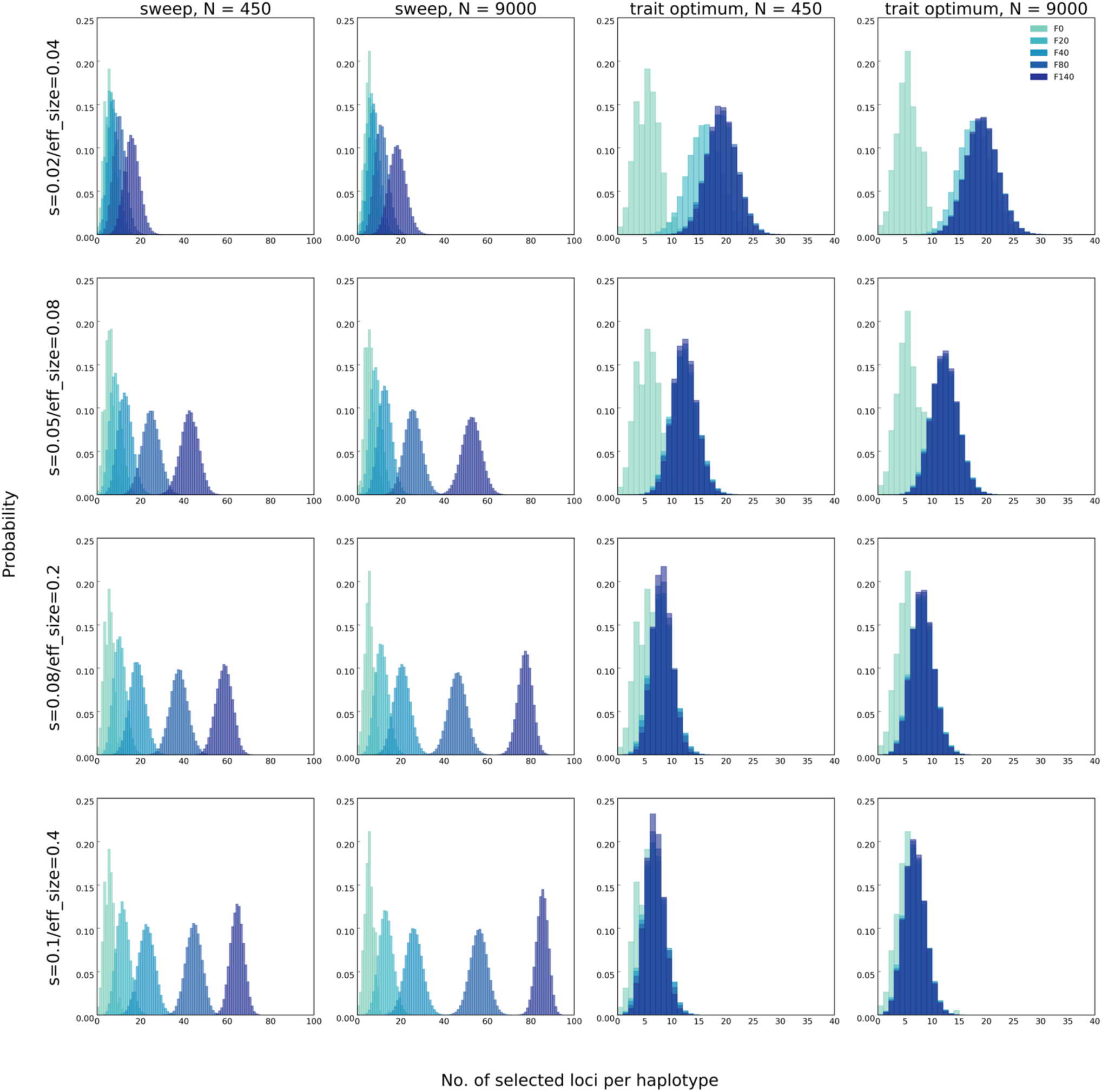
Number of beneficial loci in each haplotype in populations of 450 and 9,000 individuals under sweep and trait optimum paradigms with 100 loci and different effect sizes (scenario C in Table 1a and 1b). The area under normalized histogram equals 1. Total number of replicates in simulations are 50.

## Conclusions

Our computer simulations identified several features that can be used to distinguish between the selective sweep and trait optimum paradigms. While one distinguishing feature requires phenotypic data, majority of features can be inferred from genomic data alone provided that reaching the trait optimum is assured.

1. The fitness of large populations is greater than that of small ones under the sweep paradigm and continues to increase until the fixation of all selected alleles (Fig. 2). For the trait optimum paradigm, however, the fitness between small and large populations differs only until the trait optimum is reached and is not affected by further allele frequency changes (Fig. 2). No genetic data are required for this distinguishing feature.
2. The selected alleles increase in frequency until fixation under the sweep paradigm while the frequency of selected alleles increases until the phenotypic optimum is reached in the trait optimum paradigm (Fig. 1). After reaching the trait optimum, the median allele frequencies plateaus in large populations but not in small populations. Strong drift and frequency decrease of selected alleles below detection limit are responsible for continued increase of the median allele frequencies. Nevertheless, the number of identified selected alleles is strongly reduced in later generations in small populations. Therefore, there is a clear difference between the two paradigms in either small or large populations provided that the experiment is conducted for a sufficient number of generations.
3. The number of selected alleles shared among replicates (parallelism) is another distinguishing feature between the paradigms. The parallelism among replicates continues to increase in the sweep paradigm while after reaching the trait optimum repeatability of adaptation decreases under the trait optimum paradigm (Fig. 3). Therefore, replication provides a powerful means for distinguishing evolutionary paradigms.
4. The number of beneficial alleles per haplotype continues to increase in the sweep paradigm while it plateaus under the trait optimum paradigm (Fig. 4). This feature requires availability of phased haplotypes but provides another confirmatory test for distinguishing the two evolutionary paradigms.

As a consequence of consistent frequency increase in the sweep paradigm, many alleles reach near-fixation frequencies, i.e. sweep-like signatures (Fig. 1). But in our simulations sweep-like signatures were extremely rare for the trait optimum paradigm. Nevertheless, we caution that this is not a very reliable discriminating feature, as unequal effect sizes and few selected loci (Fig. 8) could cause sweep-like signatures in trait optimum paradigm.

We also showed that the combination of large and small replicate populations uncovers some distinctive patterns that can be further used for developing test statistics to discriminate between the two paradigms. We propose that machine learning could be a powerful approach to exploit the described features for a quantitative approach to distinguish between the two paradigms. We consider combining the analysis of small and large populations as a suitable means for the analysis of the adaptive architectures. Large populations clearly offer the advantage to identify a larger number of selected alleles which increase in frequency in multiple replicates. Small populations are easier and cheaper to maintain while still offering discriminative features. However, mapping the causative variant will be more challenging in small populations because of stronger linkage disequilibrium and more confounding signal from neutral alleles.

## Materials and Methods

We simulated a quantitative trait with linked loci under sweep and trait optimum paradigms for two population sizes, i.e. 450 and 9000 diploid individuals, assuming random mating among individuals (scenario A in Table 1a and 1b). We define the trait optimum paradigm as polygenic adaptation of a quantitative trait after a shift in phenotypic optimum. The positions of the selection targets were randomly distributed along the entire chromosomes 2 and 3 of *D. simulans*, but kept the same for sweep and trait optimum paradigms. For a realistic linkage structure and to mimic the number of haplotypes typically used in E&R studies, we used 189 haplotypes from a *D. simulans* population collected in Florida (Howie et al. 2019) to construct populations of 450 and 9000 individuals for the simulations, i.e. each haplotype is present in multiple copies in the founder population. We used the recombination landscape of *D. simulans* in our simulations (Howie et al. 2019). Population fitness (sweep paradigm) or phenotype (trait optimum paradigm) and allele frequencies were recorded every 10^th^ generation until generation 140. Each simulation scenario was performed in 500 iterations. For characterization of the qualitative differences between sweep and trait optimum paradigm we performed computer simulations using functions *w* (sweep) and *qff* (trait optimum) of MimicrEE2 (version mim2-v193) (Vlachos and Kofler 2018).

### Simulations of selective sweep paradigm

We performed forward Wright-Fisher simulations using 100 linked loci (linkage structure of the phased haplotypes (Howie et al. 2019)) with equal starting frequencies of 0.05 and equal selection coefficients of 0.08 constant across time in populations of 450 and 9000 diploid individuals for 140 generations (scenario A in Table 1a). In addition to this default scenario, we also performed simulations with different numbers of contributing loci, e.g. 10, 20, 50 and 100 (scenario B in Table 1a) and different values for the selection coefficient, e.g. 0.02, 0.05, 0.08, 0.1 (scenario C in Table 1a) in populations of 450 and 9000 diploid individuals.

### Simulations of trait optimum paradigm

In trait optimum simulations, we simulated adaptation of a quantitative trait to a new trait optimum. Trait *z* is affected by *L* diallelic loci. The effect size of the “+” allele is +*a* with frequency *p*_i_ and the effect size of the “–” allele is –*a* with frequency *q*_i_ = 1-*p*_i_. Trait *z* is computed as:

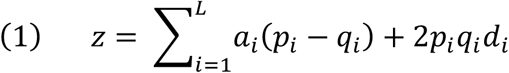

We assume co-dominance (h=0.5), *d* = 0, and epistasis is neglected, thus trait z was determined additively. The trait value is mapped to fitness (*w*) using a Gaussian fitness function where *PDF* is the probability distribution function and max*fit* and min*fit* are the maximum and minimum fitness values:

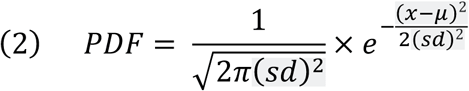

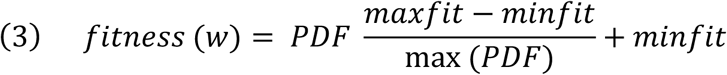

We simulated 100 linked loci starting at frequency of 0.05 with equal effects i.e. 0.04 in populations of 450 and 9000 diploid individuals (scenario A in Table 1b). The trait optimum (phenotype) was set at −2.5 (µ) with standard deviation (*sd*) of 0.3 and fitness ranged between 0.5 and 4.5 (scenario A in Table 1b). In addition, further simulations with different number of loci (scenario B in Table 1b) and different values for effect sizes (scenario C in Table 1b) were performed; for each simulation run the same effect sizes were used for all loci. The phenotypic value of the populations at the beginning of the simulations varies depending on the effect size and the number of loci. To enable comparison of simulations with different number of contributing loci and/or different effect sizes, independent of the phenotypic variance in the founder population, we adjusted the phenotypic optimum for each simulation scenario such that all populations move the same distance in the phenotypic space to reach the phenotypic optimum (Fig. S4).

### Neutral simulations

To account for the effect of drift in allele frequency changes, we performed simulations for populations with 450 and 9000 individuals with no selection; all parameters of simulations matched scenario A in Table 1a but without selection. We determined the allele frequency changes (AFC), and set the threshold for identification of alleles with AFC more than expected under drift based on the upper 5% tail of neutral AFC distribution between the founder and evolving populations at each timepoint across 500 replicates (Fig. S1).

### Repeatability of adaptation (similarity among replicates)

The average pairwise Jaccard indices (Jaccard 1901) among the replicates were calculated for 50 sets of 10-replicate populations of the sweep and trait optimum simulations using the number of alleles with allele frequency changes more than expected under drift (neutral simulations above).

## Acknowledgments

We thank Joachim Hermisson for comments on an earlier version of manuscript and Robert Kofler and Christos Vlachos for support with MimicrEE2 simulations.

