## Supplementary figures for "Distinct patterns of selective sweep and polygenic adaptation"

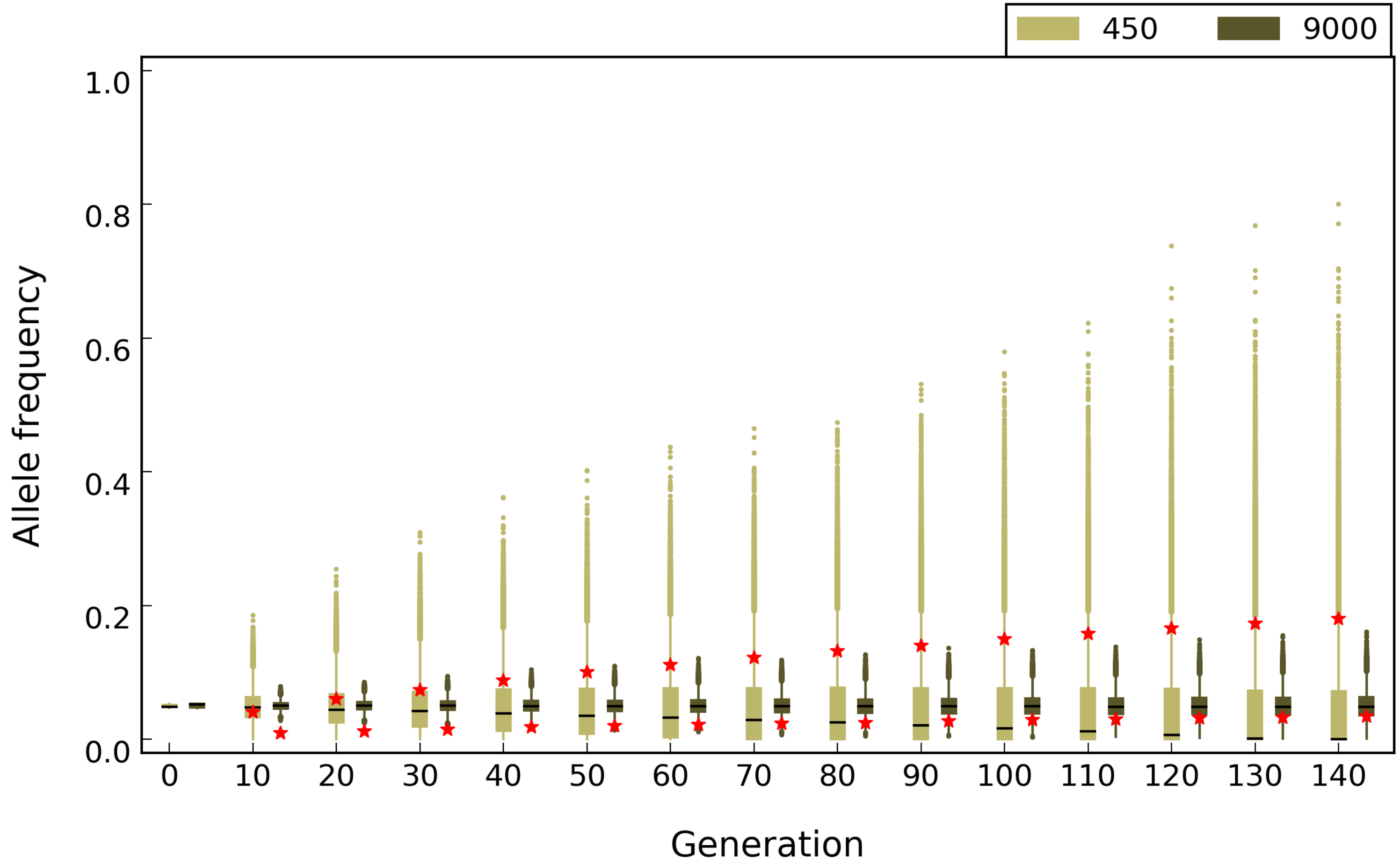


**Figure S1** Allele frequencies under neutrality. Simulation parameters were similar to scenario A (Table 1b) with no selection. Red asterisks depict the frequency cut-off based on 95% quantile of allele frequency change under neutral simulations.

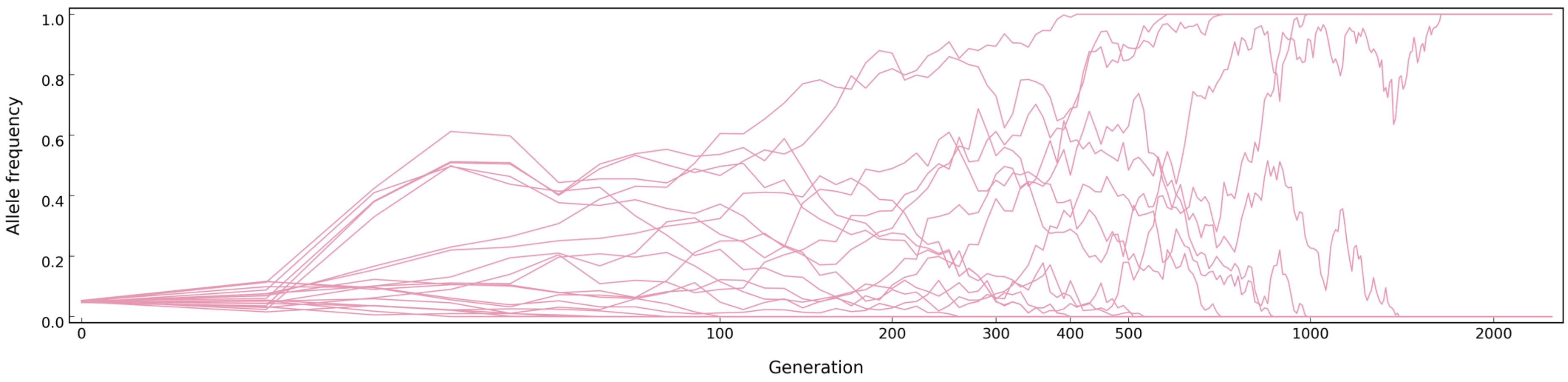


**Figure S2** Allele frequency trajectories of loci in a population of 450 individuals under the trait optimum paradigm (scenario A in Table 1b). After reaching the trait optimum at generation 30 (phase 1), the drift phase (phase 2) starts. In phase 3, all alleles are sorted, i.e. either fixed or lost. X axis is log10-transformed and for clarity only 20 out of 100 loci are shown.

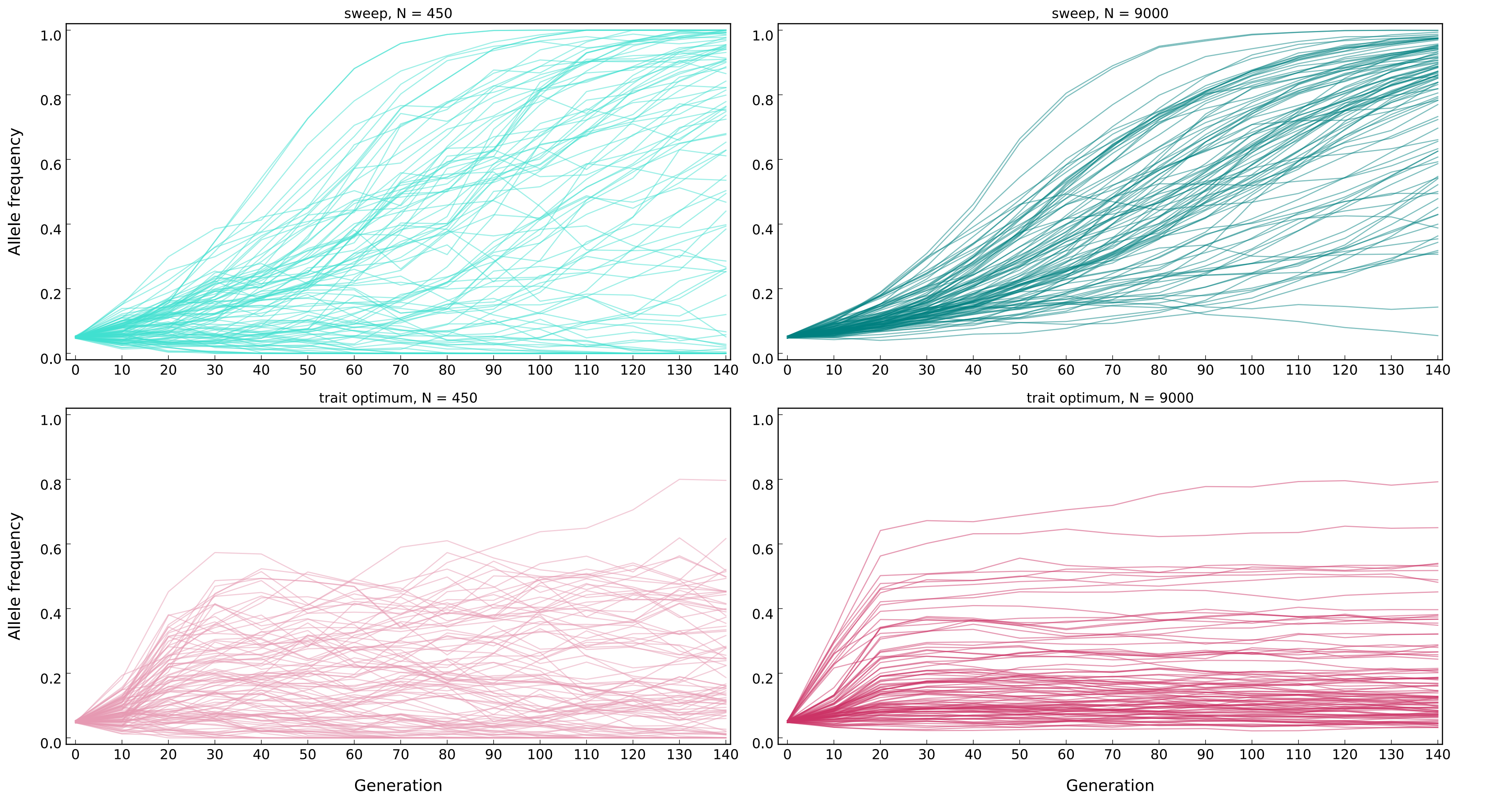


**Figure S3** Allele frequency trajectories of one simulation as an example for scenario A (Table 1a and 1b) under sweep (top panels) and trait optimum (bottom panels) paradigms.

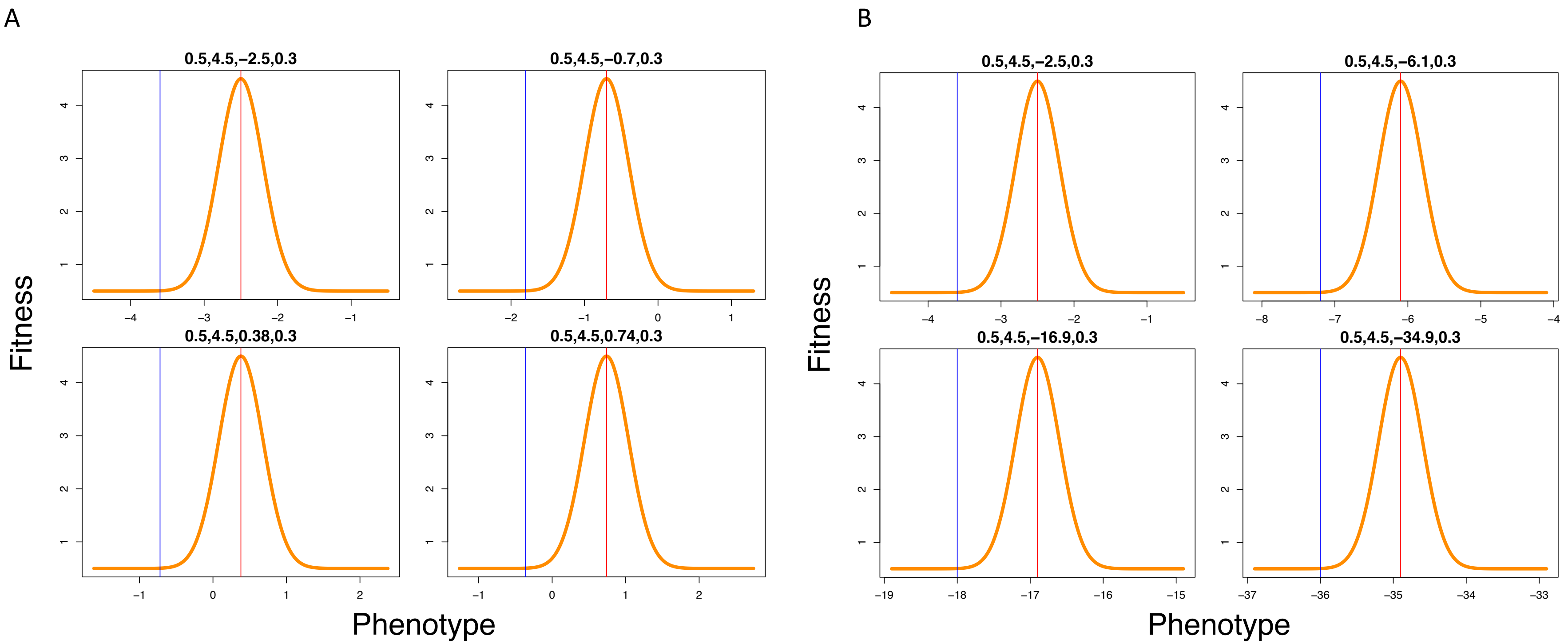


**Figure S4** Guassian fitness function for the trait optimum paradigm for A) different number of loci (scenario B in Table 1b), and B) different effect sizes (scenario C in Table 1b). The phenotype optimum is shifted so that the phenotype of the population regardless of the number of loci or their effect sizes is equally distant from the optimum. Blue line depicts the starting phenotype of the population and the red line shows the optimum phenotype. A from top to bottom (left to right) shows fitness functions for 100, 50, 20 and 10 loci, B from top to bottom (left to right) shows fitness functions for 0.04, 0.08, 0.2 and 0.4 locus effect size.
